# The Effects of Propofol Anaesthesia on Molecular-enriched Networks During Resting-state and Naturalistic Stimulation

**DOI:** 10.1101/2022.05.24.493298

**Authors:** Timothy Lawn, Daniel Martins, Owen O’Daly, Steve Williams, Matthew Howard, Ottavia Dipasquale

## Abstract

Placing a patient in a state of anaesthesia is crucial for modern surgical practice. However, the mechanisms by which anaesthetic drugs, such as propofol, impart their effects on consciousness remain poorly understood. Propofol potentiates GABAergic transmission, which purportedly has direct actions on cortex as well as indirect actions via ascending neuromodulatory systems. Functional imaging studies to date have been limited in their ability to unravel how these effects on neurotransmission impact system-level dynamics of the brain. Here, we leveraged advances in multi-modal imaging, Receptor-Enriched Analysis of functional Connectivity by Targets (REACT), to investigate how different levels of propofol-induced sedation alters neurotransmission-related functional connectivity (FC), both at rest and when individuals are exposed to naturalistic auditory stimulation. Propofol increased GABA-A- and noradrenaline transporter-enriched FC within occipital and somatosensory regions respectively. Additionally, during auditory stimulation, the network related to the vesicular acetylcholine transporter showed reduced FC within the right superior temporal gyrus, regardless of level of anaesthesia, and a spatial configuration correlating with a broad range of meta-analytic measures of audition- and emotion-related cognition. In bringing together these micro- and macro-scale systems, we provide support for both direct GABAergic and indirect noradrenergic-related network changes under anaesthesia and describe a cognition-related reconfiguration of the cholinergic network, highlighting the utility of REACT to explore the molecular substrates consciousness and cognition.

## Introduction

The ability of anaesthesia to transiently and reversibly disrupt conscious experience has both revolutionised modern surgical practice as well as provided a unique opportunity to link consciousness to its neurobiological substrates. However, the mechanisms through which this altered state of consciousness emerges are far from fully elucidated, in part due to the multiplicity of contributing systems which interact at multiple levels (Alkire, Hudetz and Tononi, 2008; Franks, 2008; Brown, Lydic and Schiff, 2010). A thorough characterisation of these different systems supporting consciousness may also offer novel therapeutic targets for disorders of consciousness, for which we remain largely bereft of meaningful treatments. As such, mechanistic investigation of anaesthetic agents and their pharmacodynamic effects may offer fundamental insights into the brain, with applications to both health and disease.

Much neuroimaging work investigating anaesthesia to date has leveraged the powerful analytic approaches applied to blood-oxygen level dependent (BOLD) signal fluctuations measured with functional magnetic resonance imaging (fMRI) whilst participants are at rest. Far more than a baseline, this resting activity reflects inherent functional organisation of the brain as well as personal mentation (Binder *et al*., 1999; Stark and Squire, 2001; Buckner, 2012). Anaesthesia seems to preferentially perturb certain domains of brain function, having been described to reduce within and between higher-level network connectivity, while preserving sensory processing within lower-level primary sensory cortices (Boveroux *et al*., 2010; Mhuircheartaigh *et al*., 2010; Stamatakis *et al*., 2010; Schrouff *et al*., 2011; Schröter *et al*., 2012; Gili *et al*., 2013; Gómez *et al*., 2013; Guldenmund *et al*., 2013; Monti *et al*., 2013; Adapa *et al*., 2014; Tang and Ramani, 2016; Liu *et al*., 2017; Qiu *et al*., 2017; Naci *et al*., 2018; Nir *et al*., 2019; Craig *et al*., 2021; Spindler *et al*., 2021). As such, studies considering only the resting state are limited in their ability to comprehensively investigate anaesthesia’s effects on the brain. Naturalistic stimuli offer a powerful tool to engage ecologically meaningful sensory and higher-level cognitive processes that do not also necessitate responses (Finn, 2021). Therefore, they are well suited to examine how the neural substrates of perception and cognition persist or are extinguished under anaesthesia.

The most widely used anaesthetic agents (including propofol, sevoflurane, and isoflurane) potentiate GABA-mediated inhibition, which alters activity in networks spanning brainstem, thalamic, and cortical regions (Brown, Lydic and Schiff, 2010). However, whether these network changes, and concomitant transitions into and out of consciousness, are mediated by top-down (direct modulation of cortical and thalamocortical circuits) or bottom-up (ascending sub-cortical arousal systems exerting influence over cortex) remains contentious (Mashour and Hudetz, 2017). There are a host of highly conserved brainstem, midbrain, and forebrain nuclei whose widespread innervation exerts neuromodulatory control over the rest of the brain (Marder, 2012; Avery and Krichmar, 2017; Shine *et al*., 2019). These can modulate the gain of receptive neuronal populations through altering their electrical and synaptic properties, thus also affecting subsequent downstream inter-regional communication (Aston-Jones and Cohen, 2005). Moreover, these systems can act in concert to produce an adaptive system that shapes network topologies and dynamics (Brezina, 2010). Functional integrity of the default mode network (DMN), which is associated with autonoetic consciousness (Liu *et al*., 2015; Guldenmund *et al*., 2017), is reportedly under the neuromodulatory influence of dopaminergic FC during propofol anaesthesia (Spindler *et al*., 2021). Conversely, another recent study highlighted the importance of direct action of propofol on cortical regions with networks showing reduced connectivity under anaesthesia also highly expressing parvalbumin positive GABAergic neurones (Craig *et al*., 2021). As such, both neuromodulatory and cortico-centric mechanisms seem to play a role (Lee and Mashour, 2018; Nguyen and Postnova, 2021), with a paucity of studies examining these effects in combination. Crucially, these need not be mutually exclusive, and it has been suggested that bottom-up and top-down mechanisms modulate separable dimensions of consciousness (Mashour and Hudetz, 2017).

Comprehensive accounts of brain function must integrate micro-, meso- and macro-scale mechanisms across different neural states (Bassett and Sporns, 2017). Conventional rest, task, and naturalistic fMRI analyses are inherently incapable of providing insights into the molecular underpinnings of the BOLD signal. This limits theoretical understanding by leaving a conceptual void between receptor level mechanisms and systems level dynamics. One solution to this has been to incorporate molecular information from positron emission tomography (PET) and single-photon emission computerized tomography (SPECT) into fMRI analyses, as in Receptor-Enriched Analysis of functional connectivity by targets (REACT), to help bridge the gap between these micro- and macro-scale systems (Dipasquale *et al*., 2019). The resultant receptor-enriched networks have demonstrated alterations under pharmacological challenge (Dipasquale *et al*., 2019, 2020; Lawn *et al*., 2022) and within disease states (Cercignani *et al*., 2021; Martins *et al*., 2022; Wong *et al*., 2022), but also offer a promising tool to probe the molecular substrates of cognition and consciousness. Each modulatory system engages with a set of target receptors which exhibit diversity in their patters of expression as well as downstream effects which interact in a complex pleiotropic manner. However, each system also has transporters, engaged in movement of neurotransmitters across vesicular and synaptic membranes, which serve as a coarse grain marker for innervation and influence over a given brain region. Moreover, the widespread arborisation of projections from these small nuclei produces a spatiotemporal influence over the BOLD signal that lends itself well to REACT. As such, these transporters offer a powerful means by which to derive distinct molecular networks associated with each system that can provide new insights into the effects of anaesthesia on the brain, both at rest and during sensory stimulation.

Here, we aimed to link functional differences across levels of consciousness, as modulated by propofol anaesthesia, with the underlying molecular systems, to identify the neurotransmitters which might play a central role in anaesthesia-induced functional changes in the human brain. We explored the molecular-enriched functional architecture of the brain and its changes under anaesthesia both at rest and under a naturalistic stimulation, hypothesising that propofol would produce divergent effects on connectivity of neuromodulatory systems at rest and during naturalistic stimulation. Specifically, we performed secondary analyses of an openly available fMRI data of healthy subjects collected at rest and whilst listening to an emotionally engaging story. Both conditions were acquired under three states of consciousness, dependent on the level of propofol administered: awake, light anaesthesia, deep anaesthesia. We derived receptor-enriched networks associated with the transporters of the main modulatory neurotransmitter systems, namely noradrenaline (NAT), dopamine (DAT), serotonin (SERT), and vesicular acetylcholine (VAChT), as well as the inotropic GABA-A receptor. These systems reflect bottom-up neuromodulatory as well as predominantly cortical primary pharmacological mechanisms respectively. We then assessed which of these networks demonstrate functional changes induced by the different levels of propofol anaesthesia; examined if these networks are significantly reshaped by the highly engaging external auditory drive, contextualising emerging differences between receptor-enriched networks at rest and under naturalistic stimulation by linking them to domains of cognition derived from Neurosynth; and tested if these conditions (rest and naturalistic stimulation) are differentially affected at varying levels of anaesthesia.

## Methods

### Participants

In this work, we employ data from previously published studies (Naci *et al*., 2018; Kandeepan *et al*., 2020) made publicly available on the OpenNeuro data repository (doi: 10.18112/openneuro.ds003171.v2.0.0). This includes data from 17 healthy subjects (Age: 24± 5, M/F: 13/4) who were right-handed, native English speakers, and showed no history of neurological disorders. The original study gained full ethical approval from the Health Sciences Research Ethics Board and Psychology Research Ethics Board of Western University (REB #104755).

### Study design

Participants underwent an fMRI scan whilst listening to an audio clip and then at rest during four sequential levels of consciousness: awake, light anaesthesia, deep anaesthesia, and recovery. The audio clip was a 5-minute excerpt from the movie “taken” depicting a teenage girl being kidnapped, intended to be highly emotionally evocative and arousing. Both story and resting-state runs were conducted with closed eyes for all states of anaesthesia.

### Anaesthesia

Here, we employ only data collected in the awake, light, and deep states. These were defined as 1) Awake: Prior to propofol administration, participants were fully awake, alert, and communicative. 2) Light anaesthesia: propofol infusion commenced with a target effect-site concentration of 0.6 µg/ml and oxygen titrated to maintain SpO2 above 96%. Once the baseline target effect-site concentration was achieved, participants’ level of sedation was assessed. Propofol produced increased calmness and slowed verbal responsiveness. Participants were considered lightly anaesthetised (Ramsey level 3) when they stopped engaging in spontaneous conversation, speech became sluggish, and only responded to loud commands. Once achieved, the effect-site concentration was maintained. 3) Deep anaesthesia: the target effect-site concentration was further increased in increments of 0.3 µg/ml with repeated assessments of responsiveness. Participants were considered deeply sedated (Ramsey level 5) when they stopped responding to verbal commands, were unable to converse, and were only rousable to light physical stimulation. Once reached, the participant was maintained at that level. Participants remained capable of spontaneous cardiovascular function and ventilation.

### MRI acquisition

Participants were provided with noise cancelling headphones (Sensimetrics, S14; www.sens.com) to deliver sound at an individualised volume deemed comfortable. Imaging was performed on a 3T Siemens Tim Trio system with a 32-channel head coil. Subjects underwent audio and resting state fMRI scans using the same BOLD EPI sequence for both conditions (33 slices, voxel size: 3mm^3^ isotropic, inter-slice gap of 25%, TR = 2000ms, TE = 30ms, matrix size = 64×64, FA = 75°). The audio and resting-state scans had 155 and 256 volumes respectively. Anatomical scans were also obtained using a T1-weighted 3D Magnetization Prepared - Rapid Gradient Echo (MPRAGE) sequence (voxel size: 1mm^3^ isotropic, TR = 2.3, TE = 4.25 ms, matrix size = 240 × 256 × 192, FA = 9°).

### Image pre-processing

Data was pre-processed using FMRIB Software Library (FSL) (https://fsl.fmrib.ox.ac.uk/fsl/fslwiki/). The processing steps included volume re-alignment with MCFLIRT (Jenkinson *et al*., 2002), non-brain tissue removal utilising the brain extraction tool (BET)(Smith, 2002), spatial smoothing with a 6mm FWHM Gaussian Kernel, and denoising utilising the Independent Components Analysis-based Automatic Removal Of Motion Artefacts (ICA-AROMA)(Pruim *et al*., 2015). Furthermore, subject-specific white matter (WM) and cerebrospinal fluid (CSF) masks were generated from segmentation of structural images, eroded to reduce partial volume effects with grey matter (GM), co-registered to the subject-specific functional space and used to extract and regress out mean WM and CSF signals from each participants functional images. Finally, data was high-pass temporal filtered with a cut off frequency of 0.005 Hz, normalised to the standard MNI152 template space, and resampled at 2 mm^3^ resolution.

### Population-based molecular templates

We employed transporter and receptor density maps from the noradrenergic, dopaminergic, serotonergic, cholinergic, and GABAergic systems (fig.1). The NAT template was derived from 10 healthy individuals utilising S, S-[^11^C]O-methylreboxetine PET (Hesse *et al*., 2017). DAT is from a publicly available template of 123I-Ioflupane single-photon emission computerized tomography (SPECT) images from 30 healthy subjects (HS) without evidence of nigrostriatal degeneration (https://www.nitrc.org/projects/spmtemplates)(García-Gómez *et al*., 2013). The SERT map was derived from the [^11^C]DASB PET images of 16 healthy controls (internal PET database). ^18^F-fluoroethoxybenzovesamicol PET was used to produce the VAChT template from 12 healthy participants (Aghourian *et al*., 2017). The GABA-A template was derived from 6 healthy individuals utilising [^11^C]flumazenil ((Myers *et al*., 2012) as described in (Dukart *et al*., 2018)). For each template, voxels within regions used as a reference for quantification of the molecular data in the kinetic model were replaced with the minimum value across all GM voxels, in order to minimise the contribution of those regions without excluding them from the main analysis (occipital cortex for NAT and DAT as well as cerebellum for SERT and VAChT). Finally, all templates were normalised by scaling image values between 0 and 1 whilst preserving the intensity distribution.

**Figure-1:**
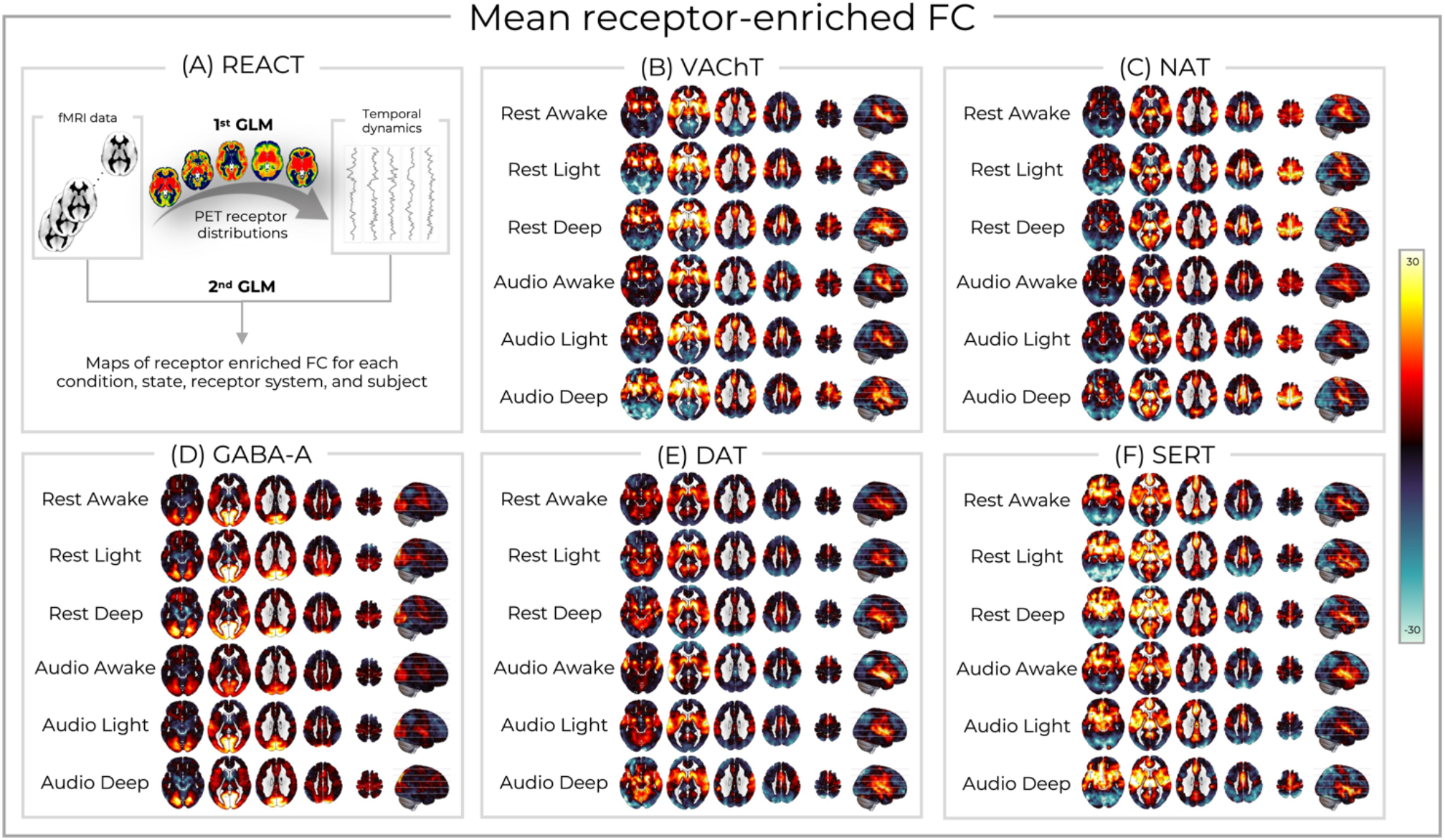
(A) The REACT analysis framework and molecular-enriched functional networks. (A) The first general linear model (GLM) extracts the dominant BOLD fluctuations within each of the PET receptor templates. The second GLM regresses these against the time series from each voxel to generate receptor-enriched maps of FC associated with VAChT (B), NAT (C), GABA-A (D), DAT (E), and SERT (F). These networks are shown averaged across participants for each condition (rest/audio) and state (awake/light anaesthesia/deep anaesthesia).

### Receptor-enriched analysis of functional connectivity

The functional networks enriched by the molecular systems (NAT, DAT, SERT, VAChT, and GABA-A) were estimated for each condition (audio and rest) and state (awake, light anaesthesia, and deep anaesthesia) using a two-step multivariate regression analysis (Dipasquale *et al*., 2019) implemented in the REACT toolbox (https://github.com/ottaviadipasquale/react-fmri (Dipasquale and Frigo, 2021)). The molecular templates were used in the first regression analysis as spatial regressors to estimate the dominant BOLD fluctuation of the functional network related to each molecular system. At this stage, both the fMRI data and the design matrix (i.e., the molecular templates) were demeaned and masked using a binarized GM atlas derived from all the molecular data, to restrict the analysis to only those GM voxels for which receptor density information was available. The resulting subject-specific time series were then used as temporal regressors in the second multivariate regression analysis, to estimate the subject-specific target-enriched functional maps. This second step was restricted to a binarized GM mask derived from all participants. Again, both data and design matrix (i.e., the time series estimated in the first step) were demeaned, with the latter also being normalised to unit standard deviation. The receptor-enriched network maps were averaged across participants for each combination of condition and state for visualisation purposes. The networks derived at rest in the awake state were also further anatomically contextualised by calculating the probability of them including each region in the Harvard-Oxford cortical and sub-cortical atlases (SI fig-1). These values were determined by thresholding the mean molecular-enriched FC maps arbitrarily at 3 to derive the rough network configuration (these images were not used for any statistical inference) before using the FSL “atlasquery” command to anatomically label regions belonging to each network.

### Statistical analysis

A repeated measures ANOVA was implemented within the Multivariate and Repeated Measures (MRM) toolbox (McFarquhar *et al*., 2016) to compare networks across conditions and states. For each receptor system, we ran a 2 × 3 repeated measures ANOVA with the within subject factors condition (rest/audio) and state (awake/light anaesthesia/deep anaesthesia). Each model used 5000 permutations and cluster-based thresholding (cluster-forming threshold *p* = 0.001). Results were family wise error FWE corrected for multiple comparisons as well as Bonferroni corrected across receptor systems (p < 0.05 / 5). Mean receptor-enriched FC was extracted from significant clusters for each participant, condition, and state. These were used to compute lower-level post-hoc pairwise tests with Bonferroni adjustment within SPSS (version 27) in order to determine which states were driving the significant ANOVA results.

Additionally, to help contextualise the results of the comparison between conditions (rest/audio), the networks showing significant differences between rest and audio were averaged across participants separately for the rest and auditory conditions and tested for relationships to domains of cognition in the form of meta-analytic maps extracted from Neurosynth. Neurosynth is an automated meta-analytic tool that integrates results from over 15,000 fMRI studies to link key search terms to voxel-wise results (https://github.com/neurosynth/neurosynth). Here, we employ a select subset of terms relevant to our resting and auditory conditions from a more comprehensive list of neurocognitive terms previously employed in conjunction with Neurosynth (Alexander-Bloch *et al*., 2018; Hansen *et al*., 2022). An in-house MATLAB script was used to estimate the Pearson’s correlation coefficient between the receptor-enriched FC maps and each cognitive term, with a spin test to provide a *p*-value corrected for spatial autocorrelation.

## Results

The REACT analysis delineated receptor-enriched FC maps associated with each neurotransmitter system for each participant, state, and condition. Averaged across participants, these molecular-enriched functional systems showed overlapping yet distinct patterns of connectivity between receptor systems as well as some degree of modification across conditions and states (fig-1). The networks from the resting awake state were further contextualised through quantifying the probability of each anatomical region being included within each molecular-enriched system (SI fig-1). Positive receptor-enriched FC of varying strength was seen within similar bilateral temporal, opercular, and insular regions across all the modulatory systems, highlighting a potential focal region for neuromodulatory molecular-enriched networks. The SERT network showed additional hubs within the frontal pole, anterior cingulate, and paracingulate cortex. NAT-enriched FC was particularly strong within the pre- and post-central gyri. Both the DAT and VAChT networks showed strong connectivity within the caudate and putamen, though the former was stronger, on average, within the lingual gyrus, precuneus, and brainstem whilst the latter was stronger within the anterior cingulate, paracingulate, and thalamus. Finally, the GABA-A network showed primary hubs within the occipital pole and lingual gyrus, but also the intracalcarine, lateral occipital, cuneus, and precuneus cortex. For full details, see supplementary figure 1.

### Main effect of state

Altered states of consciousness were associated with changes in NAT-enriched FC within the left primary somatosensory cortex (*p* = 0.006, cluster size = 238, peak MNI [*x* = -29, *y* = -33, *z* = 64])(fig-2A/C). The post-hoc test revealed that, compared to the awake state, FC was significantly greater during light anaesthesia (mean difference = 9.07, SE =1.89, *p* < 0.001, CI = 4.01 to 14.1) and deep anaesthesia (mean difference = 12.6, SE = 3.53, *p* = 0.007, CI = 3.22 to 22.1). Similarly, GABA-A-enriched FC showed a main effect of anaesthesia within large swathes of occipital cortex (*p* = 0.001, cluster size = 778, peak MNI [*x* = -26, *y* = -78, *z* = 21])(fig-2B/D). This was also driven by increased in FC during the lightly anaesthetised (mean difference = 10.0, SE = 2.37, *p* = 0.002, CI = 3.67 to 16.3) and deeply anaesthetised (mean difference = 15.6, SE = 2.83, *p* < 0.001, CI = 8.01 to 23.1) states compared to when participants were awake. Other networks showed significant differences across states, but did not survive Bonferroni correction across systems. Specifically, these were DAT-enriched network within the right cerebellar hemispheric lobule V and VAChT-enriched network within the bilateral caudate, bilateral putamen, and right angular gyrus (SI fig-2/3).

**Figure-2:**
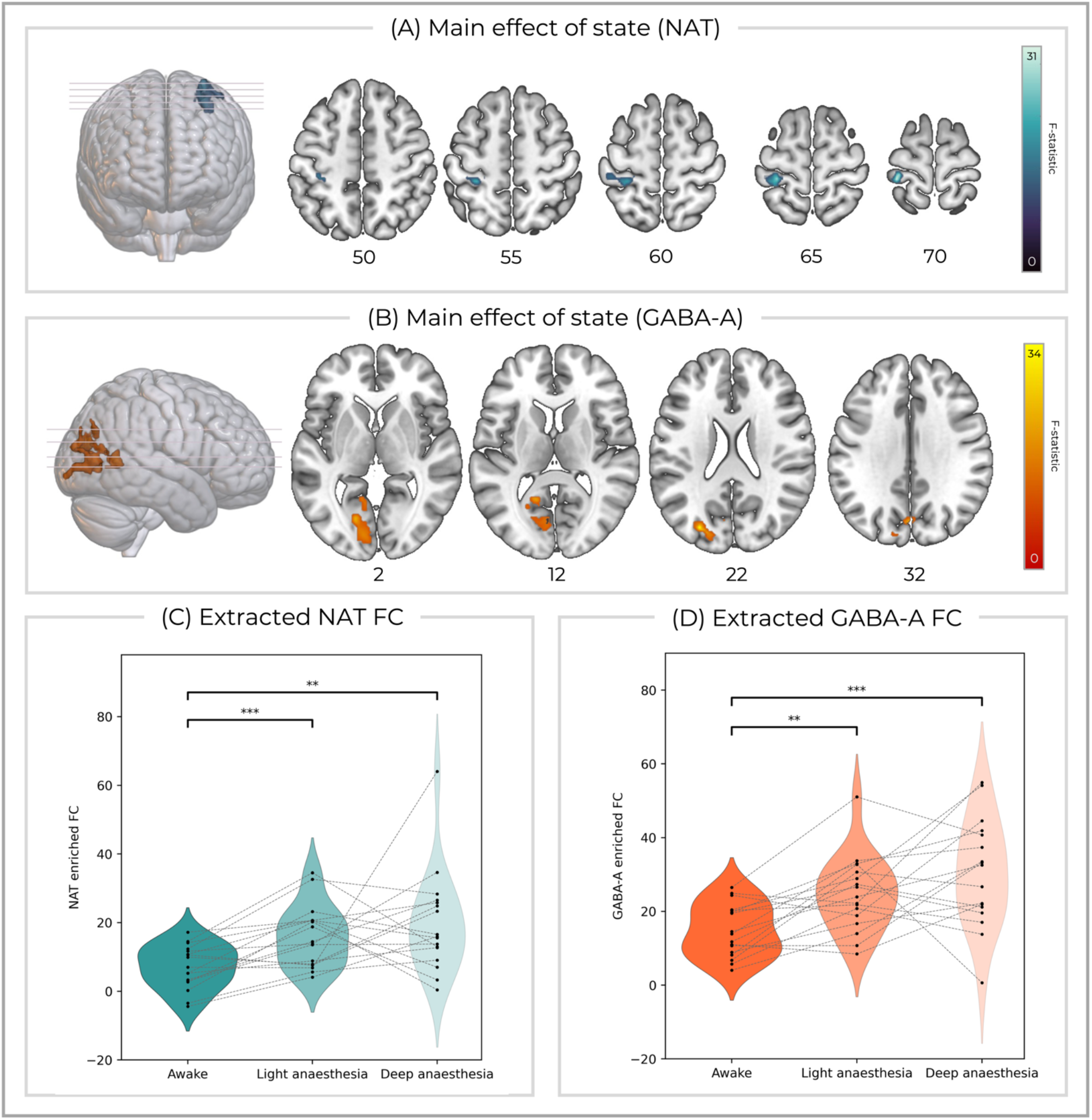
Clusters showing main effects of state (awake/light anaesthesia/deep anaesthesia) for (A) NAT-enriched FC and (B) GABA-A enriched FC that survived Bonferroni correction for multiple comparisons across systems. The receptor-enriched FC extracted from the significant (C) NAT and (D) GABA-A clusters, averaged across conditions, showed significant differences in the post-hoc contrasts [awake < light anaesthesia] and [awake < deep anaesthesia] (*** *p* < 0.001; ** *p* < 0.01).

### Main effect of condition

A significant difference between conditions of rest and auditory stimulation was found for VAChT-enriched FC, specifically located in the right superior temporal gyrus (*p* = 0.002, cluster size = 243, peak MNI [*x* = 44, *y* = -31, *z* = 13])(fig-3A). The post-hoc comparison revealed that this was driven by reduced VAChT-enriched FC during the auditory condition compared to rest (mean difference = 10.7, SE = 1.33, *p* < 0.001, CI = 7.90 to 13.5)(fig-3B). Post-hoc correlations between the spatial distributions of VAChT-enriched FC with Neurosynth meta-analytic maps of cognitive terms showed differences between auditory stimulation compared to rest (fig-3C). Specifically, auditory stimulation was associated with a broader selection of terms related auditory processing and emotion. DAT- and GABA-enriched-FC also showed a significant main effect of condition in the right middle/superior temporal gyrus and in the lateral occipital cortex respectively (SI fig-4/5), but these did not survive Bonferroni correction for multiple comparisons across systems. No significant differences between conditions were found in the NAT- and SERT-enriched functional networks.

**Figure-3:**
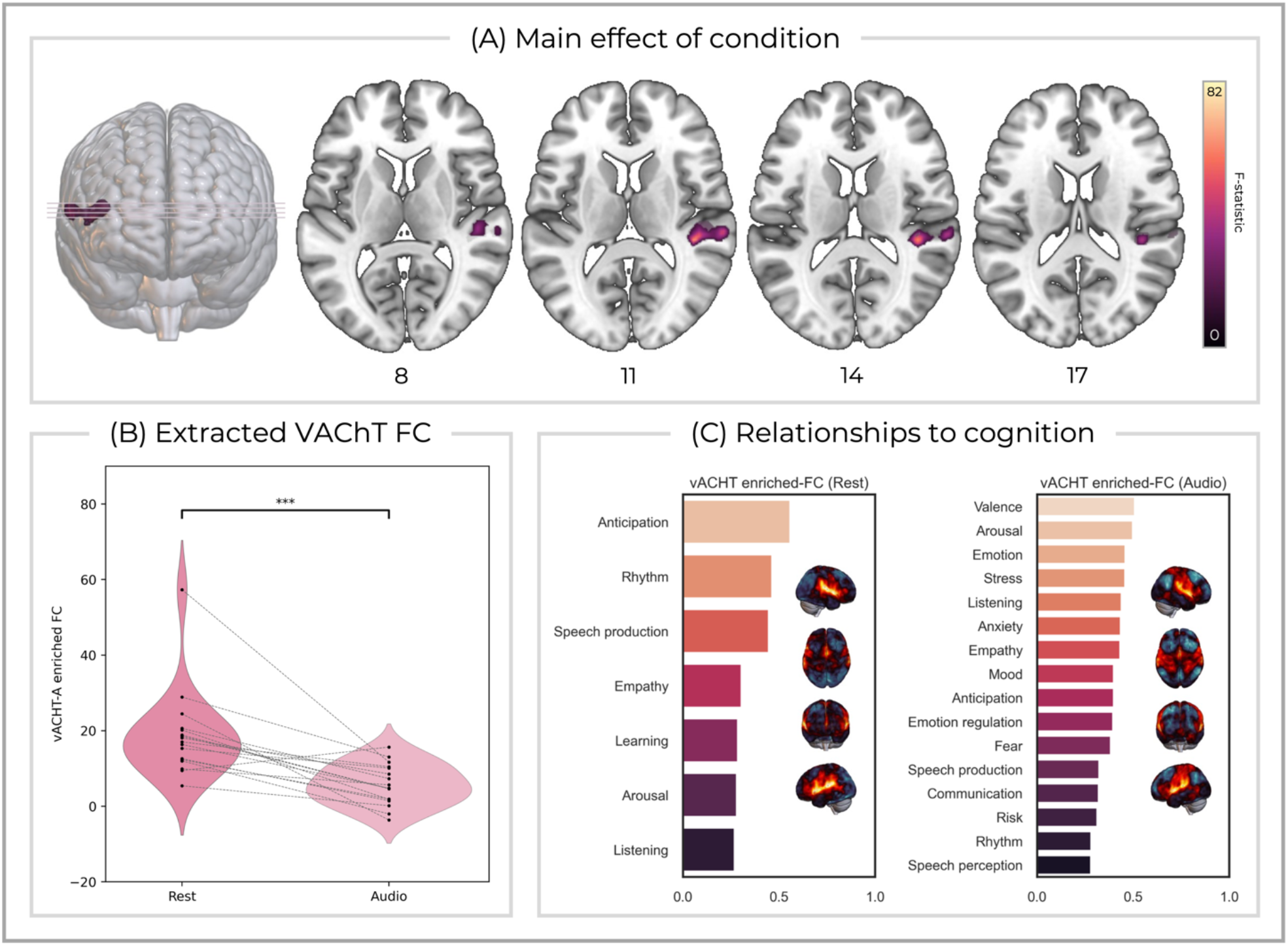
(A) VAChT-enriched FC within the right superior temporal gyrus showed a main effect of condition (rest/audio) that remained significant after Bonferroni correction for multiple comparisons across systems. The *z* MNI co-ordinate is reported below each axial slice. (B) Mean VAChT-enriched FC extracted from the significant cluster located in the right temporal gyrus, averaged across states, and displayed between conditions for each participant. (C) Significant correlation strength between VAChT-enriched FC map averaged across participants and meta-analytic results of select domains of cognition from Neurosynth for the rest condition (left) as well as naturalistic auditory stimulation (right). Each correlation is statistically significant after spin test correction for spatial autocorrelation. The VAChT-enriched FC maps are averaged across participants and 3D rendered (right, superior, anterior, and left surfaces are shown from top to bottom).

### Interaction of condition and task

We found an interaction effect in the DAT-enriched network, with the significant cluster located in the left anterior cingulate cortex (ACC) (*p* = 0.032, cluster size = 74, peak MNI [*x* = -10, *y* = 49, *z* = 4])(SI fig-6). However, this result did not survive Bonferroni correction for multiple systems. No significant interaction effects were found in the other molecular-enriched networks.

## Discussion

In this work, we explored how different levels of propofol anaesthesia shape the network architecture of the resting and engaged brain utilising novel multi-modal methods which establish clearer mechanistic links between neurotransmission and connectivity. We enriched BOLD fMRI analysis with the distribution density of inhibitory (GABA-A) as well as modulatory (NAT, DAT, SERT, and VAChT) neurotransmitters and assessed the connectivity of these networks under propofol mediated manipulations of consciousness. Given the potential benefit of using naturalistic stimuli to engage sensory and higher-level cognitive processes (Finn, 2021), we also tested if these molecular-enriched functional networks undergo substantial reconfiguration as compared to the resting state. Our findings showed a FC increase in the GABA-A and NAT-enriched networks induced by anaesthesia within occipital and somatosensory regions respectively. We also found a significant modification of the VAChT-enriched network under external auditory drive mainly located in the right superior temporal gyrus, whose FC was significantly reduced during auditory stimulation as compared to rest, regardless of level of anaesthesia. The broader reconfiguration of this cholinergic network was associated with different patterns of correlations between VAChT-enriched FC and Neurosynth meta-analytic terms congruent with a functional role in auditory processing of an emotionally evocative story. We discuss these two main findings below.

### NAT and GABA-A enriched FC increase under propofol anaesthesia

Consciousness appears to be dependent on the correlation and anti-correlation of large-scale brain networks (Dehaene and Changeux, 2011; Dehaene *et al*., 2014; di Perri *et al*., 2016). Increasing evidence supports the notion that these networks not only depend on their stable structural connectivity, but are also shaped by a host of neuromodulatory systems that exert widespread influence over diverse but overlapping cortical and subcortical regions (van den Brink, Pfeffer and Donner, 2019). Our findings expand these accounts by showing that FC increases with depth of anaesthesia for NAT and GABA-A enriched networks within somatosensory and occipital regions, respectively. At a broader network level, these results align with the known actions of propofol directly onto GABAergic cortical circuits as well as via ascending modulatory arousal systems.

Propofol acts primarily through potentiating GABAergic transmission throughout the central nervous system (Bai *et al*., 1999; Hapfelmeier, Schneck and Kochs, 2001; Hemmings *et al*., 2005, 2019). Despite a significant number of neuroimaging studies, a comprehensive account of the relationships between this GABAergic potentiation, neural activity/connectivity, and consciousness remains elusive (Bonhomme *et al*., 2019). Even in the first PET studies on anaesthesia in humans, Alkire and colleagues speculated that regional reductions in glucose metabolic rates (which was greater within cortical than subcortical regions) may be driven by the distribution of GABA-A receptors (Alkire *et al*., 1995). Accordingly, we found that propofol anaesthesia increased FC in the functional network related to GABA-A within occipital regions, which likely reflects direct actions of propofol on cortical GABA-A receptors modulating the BOLD activity of those areas. This aligns with a recent study exploiting genomic data from the Allen Human Brain Atlas (AHBA) that demonstrated that networks with significantly reduced connectivity under propofol also show a high density of parvalbumin expressing GABAergic neurones (Craig *et al*., 2021). Their complementary approach provides cellular meso-scale insight, further linking global connectivity measures to the GABAergic system by particularly implicating this subpopulation of inhibitory neurones. Future work examining both receptor and cellular systems in combination may allow for more comprehensive mapping of the functional contributions of these lower-level organisational principals onto systems levels dynamics.

Interestingly, Craig et al. also found that many areas, including the somatosensory cortex, exhibited increases in connectivity as a function of depth of anaesthesia (Craig *et al*., 2021). Unlike the regions of reduced connectivity mentioned above, these increases were not significantly associated with GABAergic expression, alluding to an alternative mechanism. Our results demonstrate that noradrenergic FC is increased in the left somatosensory cortex during propofol sedation, suggesting that altered noradrenergic tone may contribute to these additional changes in connectivity. It is worth mentioning that although our Bonferroni corrected results are lateralised, contralateral clusters are present below significance level, indicative of a bilateral effect. Propofol has been reported to supress activity within brainstem nuclei relating to arousal (Nguyen and Postnova, 2021), including the locus coeruleus (LC)(Chen, Yang and Chiu, 1999; Du *et al*., 2018). The LC provides widespread noradrenergic projections to virtually all regions of the brain (Sara, 2009). We found that NAT-enriched FC in the somatosensory cortex increased with loss of consciousness. This may reflect suppression of LC activity by propofol, resulting in the selective modulation of BOLD signal in noradrenaline recipient regions. Furthermore, neuromodulators including noradrenaline also tune thalamocortical network synchronisation (Dahl, Mather and Werkle-Bergner, 2022), which has been strongly implicated in mechanisms of anaesthesia (Malekmohammadi *et al*., 2019). This is also concordant with various supporting evidence for a causal contribution of noradrenaline to maintaining consciousness. Dexmedetomidine, which inhibits noradrenergic neurones within the LC through presynaptic α2 adrenoceptor agonism (Nelson *et al*., 2003), produces a state similar to non-REM sleep and reduces the dose of propofol required to induce loss of consciousness (Peden *et al*., 2001; Zhang *et al*., 2021). Similarly, chemogenetic activation of NA populations within the LC can retard anaesthetic induction as well as produce cortical arousal and expedited behavioural emergence from unconsciousness following isoflurane anaesthesia in a manner preventable by α1 or ß receptor antagonism (Vazey and Aston-Jones, 2014). Furthermore, mutations perturbing NA biosynthesis can produce hypersensitivity to anaesthetic induction and particularly diminish emergence from anaesthesia (Hu *et al*., 2012). However, neither pharmacological blockade of noradrenergic reuptake (Kenny et al., 2015), nor microdialysis of noradrenaline within the prefrontal cortex (Pal *et al*., 2018) restore consciousness during continuous sevoflurane anaesthesia. Moreover, manipulations of other modulatory neurotransmitter systems including acetylcholine, dopamine, histamine, and orexin can also modulate the neurophysiological induction, maintenance, and emergence from anaesthesia (Hemmings *et al*., 2019). Thus, whilst noradrenergic transmission from the LC is clearly involved, its alteration under anaesthesia seems to be neither necessary nor sufficient for the resultant behavioural manifestation. This further highlights the complex contribution of numerous mechanisms which interact at multiple levels to enact general anaesthetic agents’ effects on consciousness.

It remains unclear why we did not identify differences in the other modulatory receptor-enriched networks under anaesthesia. In particular, projections from the dopaminergic ventral tegmental area (VTA) to the posterior cingulate cortex (PCC) and precuneus have recently been described to modulate DMN connectivity under propofol anaesthesia (Spindler *et al*., 2021). The authors also attempted to demonstrate a more direct causal role for dopamine in this mechanism by showing methylphenidate boosts VTA – PCC/precuneus connectivity in patients with disorders of consciousness. However, the temporal dynamics of the VTA and LC are positively correlated, with the collective activity of brainstem nuclei generally showing widespread anticorrelation with the cortex (Zhang *et al*., 2016). Moreover, methylphenidate also increases levels of noradrenaline, and the LC has recently been robustly demonstrated capacity to modulate frontal DMN regions (Oyarzabal *et al*., 2022). As such, disentangling the effects of these catecholaminergic systems on brain-wide connectivity is challenging and the aforementioned findings may be driven at least in part by the noradrenergic system. Indeed, both the PCC and precuneus showed similar levels of dopaminergic and noradrenergic transporter-enriched FC (SI fig-1). REACT allowed us to examine both DAT and NAT related FC simultaneously, and the distribution of their receptor sub-systems has previously been employed to attempt to delineate their associations to distinct patterns of connectivity induced by atomoxetine (van den Brink, Nieuwenhuis and Donner, 2018). Future work more selectively manipulating these systems may allow for more careful characterisation of their specific contributions to network changes under anaesthesia.

### Naturalistic stimulation induces a spatial reconfiguration of VAChT-enriched functional network regardless of anaesthesia

Analysis of task-based fMRI conventionally entails convolving an event or block related design with a haemodynamic response function and then identifying voxels whose BOLD activity shows temporal concordance with this predicted time series. Conversely, here we employ a temporally coarse-grained approach by calculating a measure of static receptor-enriched FC during naturalistic auditory stimulation and comparing this to the resting state condition. This comparison delineated a region of the VAChT-enriched network, namely the right superior temporal gyrus, that demonstrated reduced FC during the auditory condition as compared to rest. In other words, the BOLD time series of this cluster was less coupled to the dominant fluctuations of the functional network related to VAChT during the task condition than at rest. Interestingly, post-hoc correlation of VAChT-enriched FC maps averaged across participants with meta-analytic patterns of activity associated with various domains of cognition demonstrated correlations with a more extensive set of terms terms relating to auditory (e.g., listening) and emotional processing (e.g., valence, arousal, emotion, stress, anxiety, empathy) during the naturalistic stimulation than at rest. These findings suggest that the cholinergic network undergoes some degree of broader spatial reconfiguration under external auditory drive and that this reconfigured state is strongly engaged with cognition. Moreover, the auditory story was intended to be highly evocative and emotionally arousing. Therefore, there is a compelling symmetry to these findings, with emotive auditory subject matter that reshapes the cholinergic network resulting in strong correlations with Neurosynth meta-analytic terms pertaining to emotion and audition. Larger studies specifically linking reconfiguration of modulatory receptor-enriched networks under different tasks may demonstrate a powerful approach to link human cognition to its molecular substrates.

Cholinergic corticopetal projections and concomitant neurotransmission have long been implicated in mechanisms of memory, attention, and perception (Blokland, 1995; Hasselmo, 2006; Klinkenberg, Sambeth and Blokland, 2011; Decker and Duncan, 2020; Venkatesan *et al*., 2020). Protracted diurnal fluctuations in cholinergic tone track state shifts through active wakefulness to rest and sleep (Jones, 2005), which is mirrored in time-of-day variations in sustained attention (Riley *et al*., 2017; Decker and Duncan, 2020). At a mechanistic level, acetylcholine can increase the gain of cortical neurones recipient of feed-forward thalamic information; a process by which the slope of neural input output-relationships is modified, permitting changes in input sensitivity whilst maintaining input selectivity (Carandini and Heeger, 2012). It has also been suggested to augment coupling of neural responses to natural scenes, whilst desynchronising local neural populations and diminishing lateral intracortical transmission (Hsieh, Cruikshank and Metherate, 2000; Roberts *et al*., 2005; Silver, Shenhav and D’Esposito, 2008; Goard and Dan, 2009; Chen, Sugihara and Sur, 2015; Minces *et al*., 2017). Together, these mechanisms may increase the signal-to-noise ratio for sensory inputs on a background of ongoing cortical activity and diminish the impact of prior (top-down) knowledge relative to new (bottom-up) information (Hsieh, Cruikshank and Metherate, 2000; Yu and Dayan, 2005; Minces *et al*., 2017; Sarter and Lustig, 2019). However, the way these mechanisms manifest at a systems level remains less well characterised. For instance, pharmacologically boosting levels of acetylcholine decrease cortex wide interactions during rest, but not task, highlighting divergence of acetylcholine related functional connectivity between conditions (Pfeffer *et al*., 2021). As such, acetylcholine seems to cause desynchronisation both at the level of local field potentials and intracortical transmission as well as global functional connectivity, albeit in a potentially condition-dependent manner. Altogether, these data suggest that the reduced VAChT-enriched FC observed here may represent increased cholinergic tone during auditory stimulation, which in turn might result in desynchronisation of BOLD fluctuations within auditory cortex relative to those within its broader spatial distribution. The functional significance of the right lateralisation remains unclear, although attentional systems show a right hemispheric dominance (Coull, 1998; Corbetta and Shulman, 2002). Future work with more careful cognitive and pharmacological manipulation will be required to characterise links between these neurophysiological findings and specific facets of attention, perception, and cognition.

The absence of a significant interaction or a main effect of states for VAChT-enriched FC suggests that these cholinergic effects largely persist under the effects of propofol anaesthesia. Previous work largely demonstrates that, during the awake state, auditory stimulation produces significant activations in bilateral temporal and frontal regions, of which the former but not the latter show some level of preservation under propofol anaesthesia (Heinke *et al*., 2004; Plourde *et al*., 2006; Liu *et al*., 2012; Adapa *et al*., 2014). As such, the basic effects of cholinergic transmission in primary sensory cortices may also persist under anaesthesia, but higher-level mechanisms functionally and causally downstream still preclude the integration of lower-level information into a coherent percept (Dehaene *et al*., 2006, 2014; Dehaene and Changeux, 2011). Future larger studies specifically examining the interactions of different modulatory systems within the focal temporal regions showing positive receptor-enriched FC across modulatory systems, as well as the potential differential contributions of nicotinic and muscarinic receptors, will be important to further link these receptor-enriched networks to their precise functional roles in sensory processing of naturalistic stimuli.

### Limitations

This work is not without limitations. Firstly, the small sample size may limit our power to detect alterations within these receptor-enriched networks and interactions between conditions and states, especially alongside the stringent Bonferroni correction applied. Similarly, stronger doses of propofol may produce more substantial effects on molecular-enriched networks than those seen here. Secondly, the PET templates employed within the REACT analyses were average maps derived in separate cohorts of healthy individuals. Thus, accounting for inter-individual differences in receptor density in our sample is not possible. However, the use of independent average templates brings the advantage of permitting investigation of multiple targets without requiring the acquisition of multi-tracer data in the same subject, which is typically not feasible. Finally, the spatial distributions of molecular targets we studied here do show some level of correlation between each other. Though perhaps conservative, by including all maps within the same model we ensure that the variance of the BOLD signal is partitioned between all systems and allows for the delineation of distinct functional networks.

## Conclusion

In this work, we provide new evidence that propofol engages with both cortical and sub-cortical targets to shape the network architecture of the brain during anaesthesia. Furthermore, we delineate a novel cholinergic network which shows cognition-related reconfiguration, with no evidence that this mechanism is altered under propofol anaesthesia. This novel application of REACT highlights the significant potential of this methodology to further unravel the contribution of molecular systems to various facets of perception and cognition. Future work examining interactions with other transmitter systems, the contribution of receptor subtypes, and dynamic fluctuations in receptor-enriched FC may offer further critical insights into neuromodulatory contributions to processing of naturalistic stimuli. Furthermore, characterising receptor-enriched network changes across diverse anaesthetic agents may shed further light on the functional contributions of these molecular systems as well as aid identification of common paths to unconsciousness. In the longer term, an understanding of how molecular mechanisms shape the complex systems level dynamics from which consciousness emerges may offer novel opportunities to treat those suffering from disorders of consciousness.

## Acknowledgements of funding and conflicts of interest disclosure

All authors have no formal conflicts of interest to declare. TL is in receipt of a PhD studentship funded by the National Institute for Health Research (NIHR) Biomedical Research Centre at South London and Maudsley NHS Foundation Trust and King’s College London. DM, OOD, MAH, and OD are supported by the NIHR Biomedical Research Centre and Clinical Research Facility at South London and Maudsley NHS Foundation Trust and King’s College London. The views expressed are those of the authors and not necessarily those of the NHS, the NIHR or the Department of Health and Social Care.

