## supplementary figure 1 for "The Effects of Propofol Anaesthesia on Molecular-enriched Networks During Resting-state and Naturalistic Stimulation"

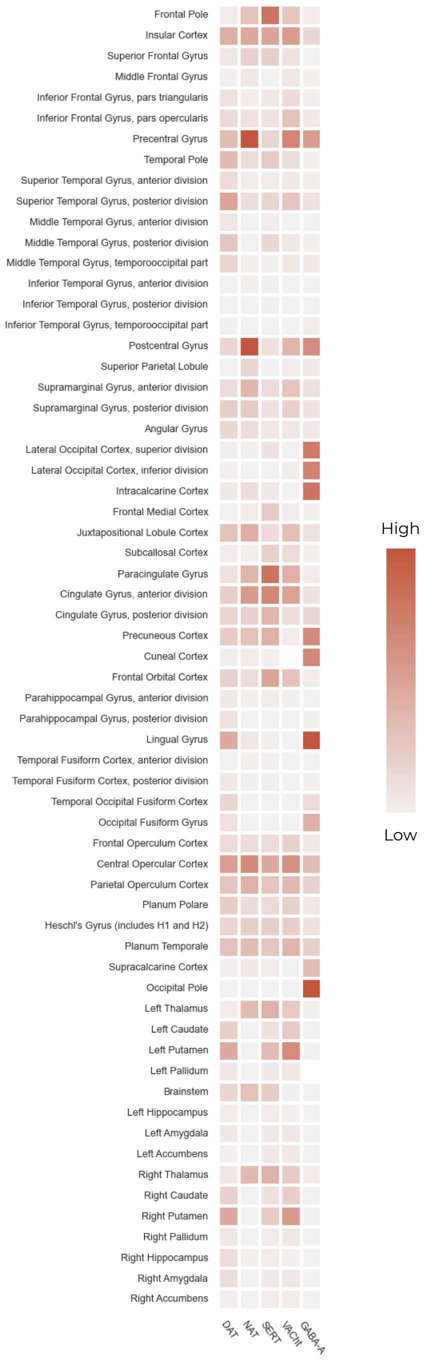


Supplementary figure 1: The probability of each anatomical region in the Harvard-Oxford cortical and sub-cortical anatomical atlases being a part of each molecular-enriched network. Values shown are only for the networks derived whilst participants were in the resting condition and awake state. These values were determined by thresholding the networks arbitrarily at 3 to derive only regions of positive FC before calling the FSL "atlasquery" command. The magnitude of the values does not convey important information, but the relative values of each network help demonstrate similarities and differences between each molecular-enriched network.

#### DAT enriched FC – Main effect of state

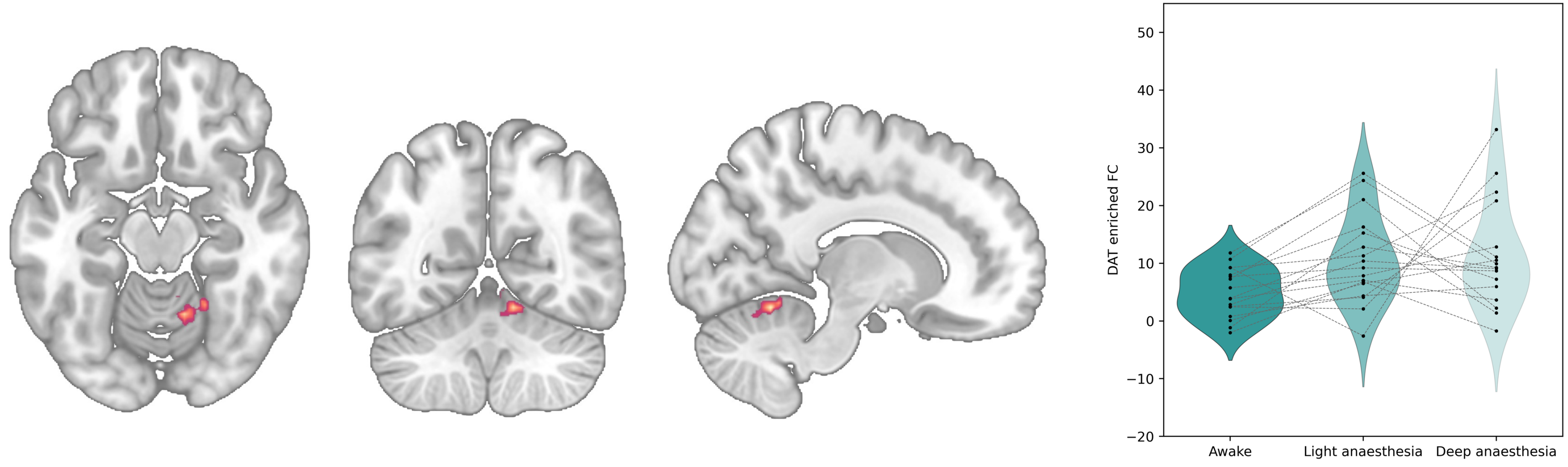


Supplementary figure 2: Cluster of DAT-enriched FC showing a significant main effect of state which did not survive Bonferroni correction across molecular systems.

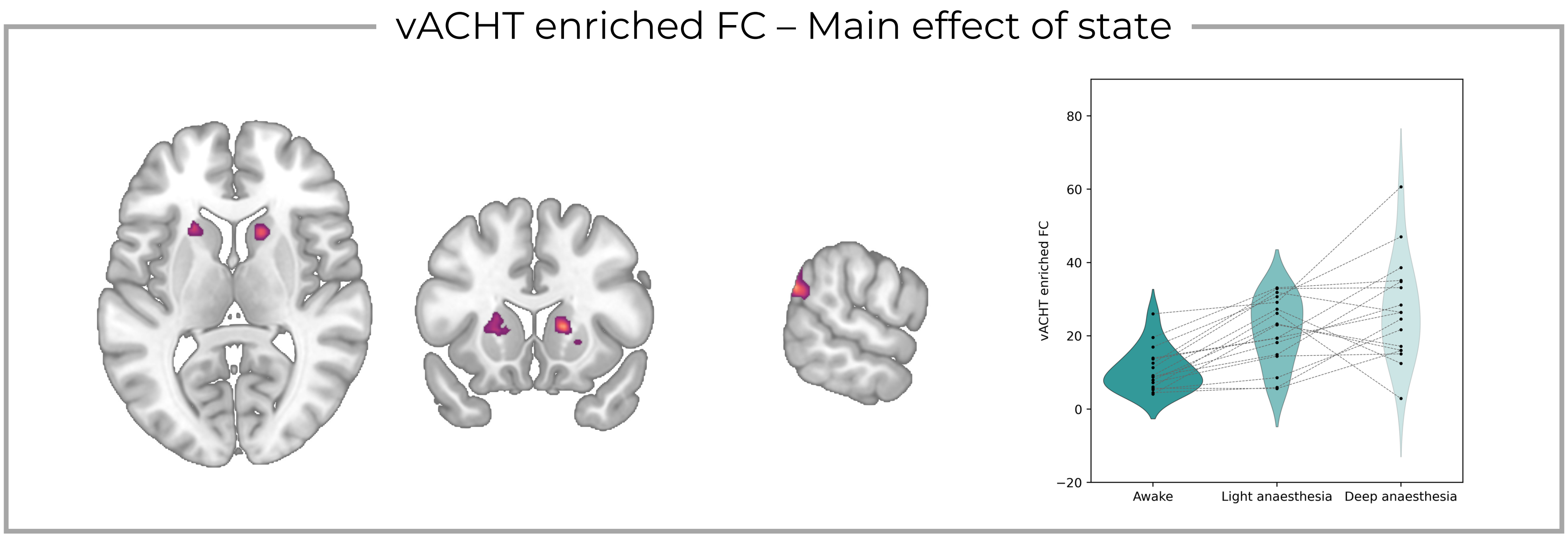


Supplementary figure 3: Cluster of VAChT-enriched FC showing a significant main effect of state which did not survive Bonferroni correction across molecular systems.

### DAT enriched FC – Main effect of condition

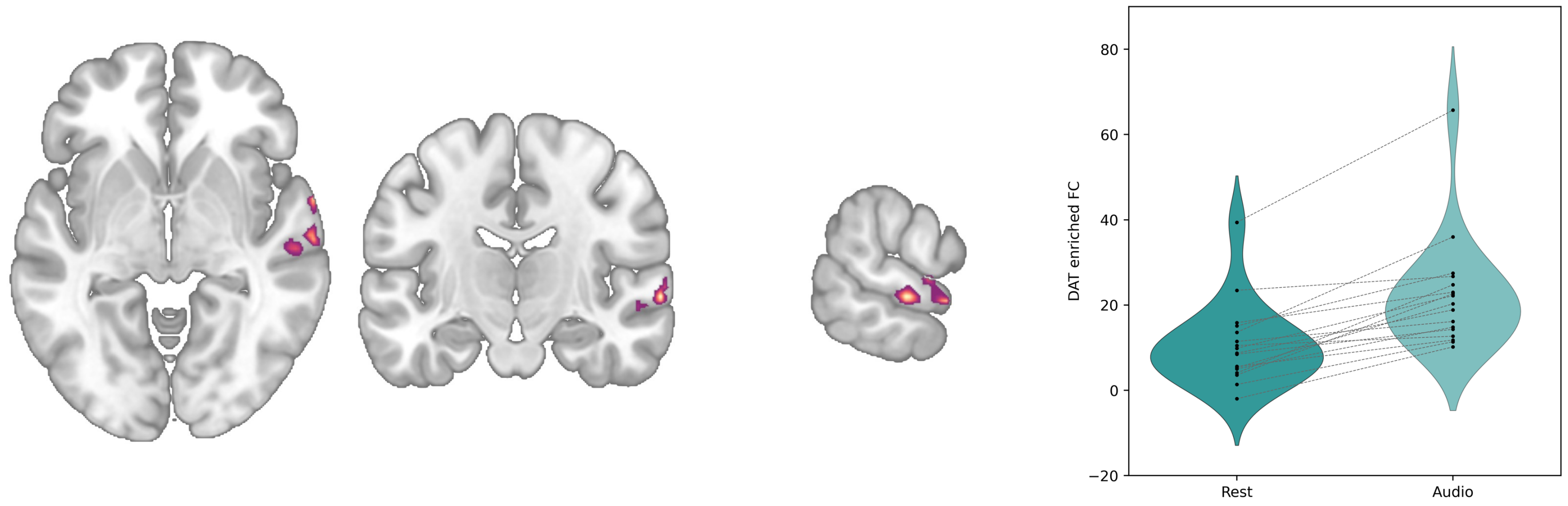


Supplementary figure 4: Cluster of DAT-enriched FC showing a significant main effect of condition which did not survive Bonferroni correction across molecular systems.

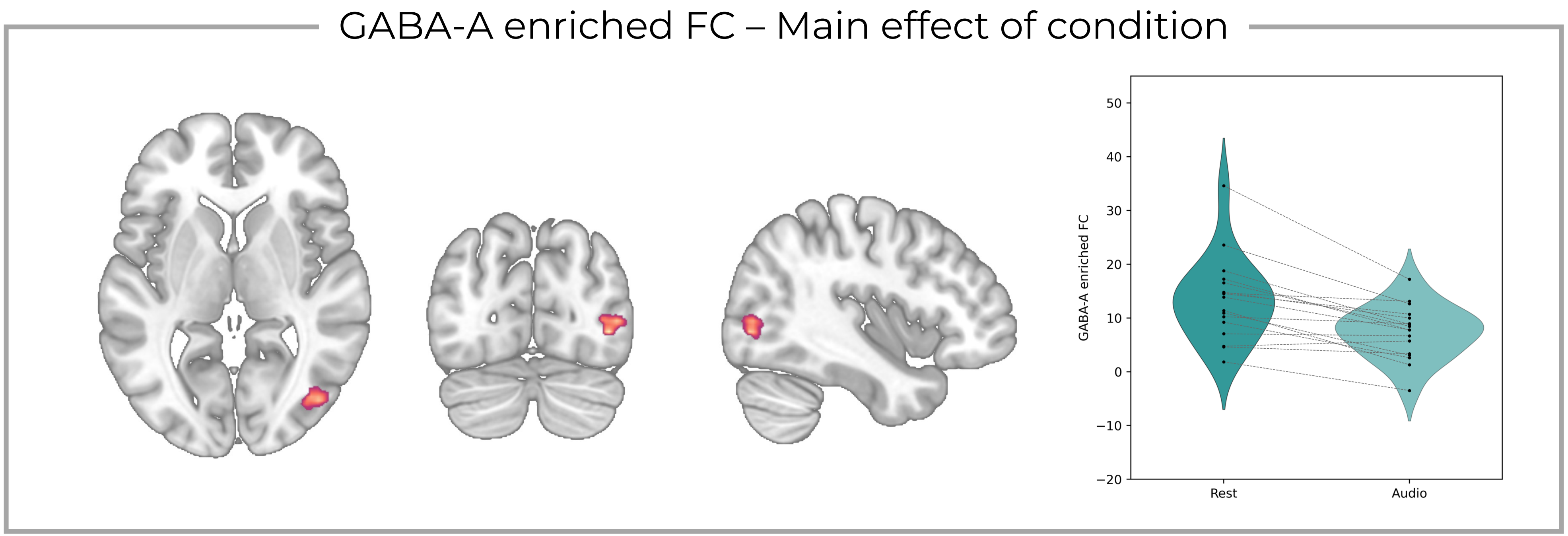


Supplementary figure 5: Cluster of GABA-A-enriched FC showing a significant main effect of condition which did not survive Bonferroni correction across molecular systems.

### DAT enriched FC – Interaction effect

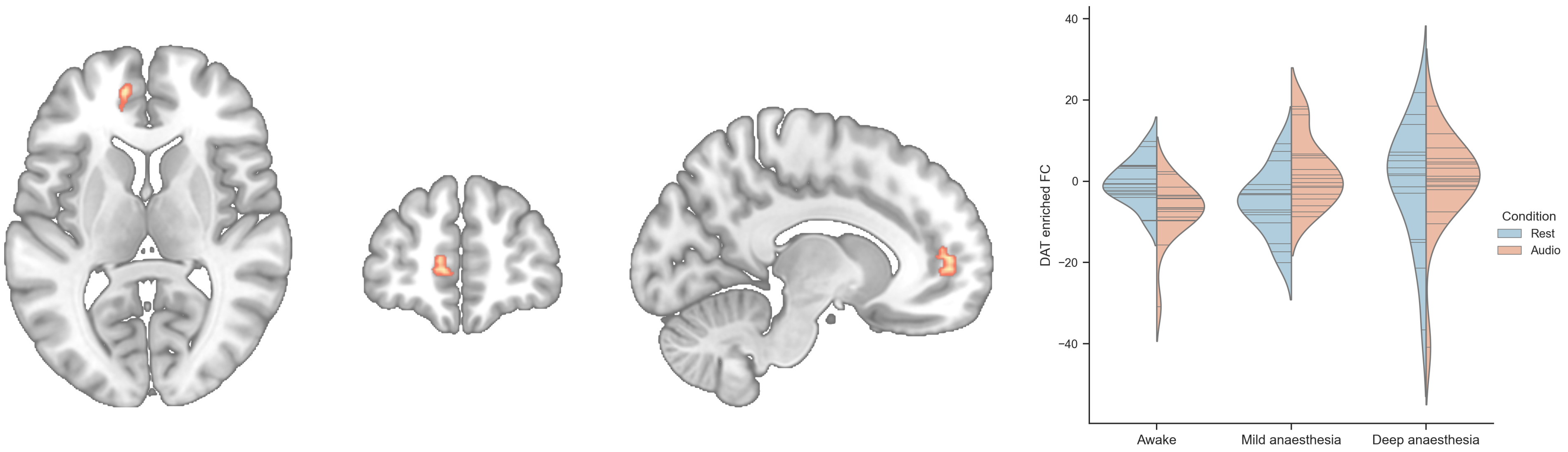


Supplementary figure 6: Cluster of DAT-enriched FC showing a significant interaction effect between condition and state which did not survive Bonferroni correction across molecular systems.
